# Subclonal IDH1/2 Mutations as a Targetable Vulnerability in Vascular Tumors

**DOI:** 10.64898/2026.05.14.725132

**Authors:** Dong-Min Yu, Eunice Lee, Gabriel J Starrett, Zili Zhai, Eleanor Dowell, Kaitlyn Walsh, Andrew Day, Doreen Palsgrove, Justin Bishop, Dylan M. Marchione, Maryam Asgari, Stephen Chung, Whitney High, Joyce MC Teng, Josh Wisell, Breelyn A. Wilky, Donald A Glass, Gregory A. Hosler, Richard C. Wang

## Abstract

Despite extensive sequencing, the genetic etiology of sporadic angiosarcoma remains poorly defined (1-3). Maffucci syndrome, characterized by vascular tumors and elevated cancer risk, is driven by mosaic gain-of-function mutations in IDH1/2 (4,5), though these have not been reported in sporadic angiosarcoma. We identify recurrent, low–variant allele frequency hotspot mutations in *IDH1/2* in over half of sporadic angiosarcomas. Mutations were validated by Sanger sequencing and immunohistochemistry. Mutant IDH1 endothelial cells promote tumorigenesis through non-cell-autonomous mechanisms, secreting 2-hydroxyglutarate (2-HG) to increase growth factor and endothelial-to-mesenchymal transition gene expression, activate pAkt/pERK signaling, induce DNA methylation changes, and promote anchorage-independent growth, which are reversed by the mutant IDH1 inhibitor ivosidenib. Patients with mosaic IDH1 mutations show reduced serum 2-HG and marked tumor regression following ivosidenib treatment. The clinical efficacy of ivosidenib in vascular tumors with subclonal IDH1 mutations suggests that low VAF *IDH1/2* mutations may be a targetable vulnerability in sporadic angiosarcoma. (6,7)

**Statement of Significance:** We identify recurrent, low-VAF IDH1/2 mutations in angiosarcoma and provide evidence that these subclonal mutations promote tumorigenesis through non-cell-autonomous mechanisms. Vascular tumors driven by subclonal IDH1 mutations responded dramatically to ivosidenib, thus revealing a novel treatment for a subset of vascular tumors.

---

Angiosarcomas are rare, frequently aggressive malignancies that arise in endothelial cells (Extended Data Fig. 1A,B; Extended Data Fig. 2). The tumor’s tendency to infiltrate normal tissues presents both diagnostic and therapeutic challenges. The tumors have a poor prognosis, with a 5-year survival of 60% with localized disease and a median overall survival of just 9-15 months with advanced disease(1-3,8). While genetic studies have provided insights into specific subtypes of angiosarcoma (e.g. secondary, primary breast), the pathogenesis of most sporadic angiosarcoma remains unclear and poses a barrier to improving the treatment of this disease.(2,9,10)

Angiosarcomas are more common in some cancer predisposition syndromes, including Maffucci syndrome—a rare, sporadic genetic disease characterized by the development of vascular neoplasms and enchondromas (Extended Data Fig. 1C,D). Vascular neoplasms, including spindle cell hemangiomas (SCH), which typically involve the skin and subcutaneous tissue, often lead to complications including pain, bleeding, and location-dependent functional impairment. Existing therapies for SCH have been limited by poor efficacy, side effects, and recurrence.(11) Maffucci syndrome is caused by mosaic, gain-of-function mutations in isocitrate dehydrogenase 1 (*IDH1*) or isocitrate dehydrogenase 2 (*IDH2*).(4,5) These mutations occur on specific residues (i.e. IDH1 R132 and IDH2 R172) and impart mutant enzymes with a neomorphic enzymatic activity. Given the role of *IDH1* mutations in the pathogenesis of the SCH of Maffucci syndrome, targeted therapies aimed at inhibiting mutant IDH1 enzymes might offer therapeutic benefits with a favorable toxicity profile but have not yet been reported.(12)

Maffucci syndrome patients are approximately 1000-fold more likely to develop a wide range of malignancies, including chondrosarcoma, glioma, cholangiocarcinoma, hematologic malignancies, and angiosarcoma.(11,13-18) When most of these cancer types occur sporadically, a proportion of the tumors also possess hotspot mutations in *IDH1/2*.(19) Curiously, sporadic angiosarcomas have not previously been reported to possess mutations in *IDH1/2*.(15,20,21) We hypothesized that a subset of sporadic angiosarcomas might possess low variant allele frequency (VAF) mutations in *IDH1/2*. Further, we propose that subclonal mutations in *IDH1/2* can promote non-cell-autonomous tumorigenic effects in angiosarcoma and other IDH1/2 mutant vascular lesions.

## Results

### Subclonal IDH1/2 mutations in angiosarcoma

To test the hypothesis that mutations in *IDH1/2* might contribute to the development of sporadic angiosarcoma, we identified archived cases of cutaneous and splenic angiosarcoma. Cancer hotspot amplicon next generation sequencing (NGS) confirmed the presence of IDH1 R132C mutations in both cases of SCH. Mutations affecting R132 in *IDH1* or R172 in *IDH2* were discovered in 77% (17/22) of the angiosarcoma. Strikingly, 27% (6/22) of angiosarcoma cases possessed multiple hotspot mutations within the same tumor (Table 1; Extended Data Table 1). The combined VAF of *IDH1/2* mutations detected by NGS ranged from 0.17-1.28%. BLAST analyses confirmed that mutations occurred on distinct alleles, and LoFreq confirmed that *IDH1/2* were significantly enriched above background mutations suggesting a convergent evolutionary process where multiple subclonal lineages independently acquire *IDH* mutations to promote tumor growth (Extended Data Table 1).(22) Amongst the non-angiosarcoma vascular lesions, a single R172K mutation was detected by amplicon sequencing in a venous malformation.

**Table 1.** IDH1 and IDH2 mutations in angiosarcoma and other vascular tumors. Summary of hotspot mutations identified by next-generation sequencing (NGS) and validated by BDA-PCR and Sanger sequencing (*n* = 25). The table lists the specific diagnoses, mutation types, and total variant allele frequency (VAF) validated across both detection platforms for each patient sample. ‘-’ BDA-PCR was not completed; ‘i.s.’ insufficient sample for BDA-PCR; ‘CA’ cutaneous angiosarcoma; ‘SA’ splenic angiosarcoma; ‘SCH’ spindle cell hemangioma; ‘LCH’ lobular capillary hemangioma; ‘AVM’ arteriovenous malformation; ‘VM’ venous malformation; ‘Total VAF’ – Only mutant alleles that were confirmed by both NGS and BDA-PCR were included in the ‘total VAF’ sum.

| Patient | Diagnosis | IDH1 R132C |  | IDH1 R132H |  | IDH2 R172K |  | IDH1/2 |
| --- | --- | --- | --- | --- | --- | --- | --- | --- |
|  |  | NGS | Sanger | NGS | Sanger | NGS | Sanger | Total VAF |
| 1 | CA1 | Yes | Yes | Yes | Yes | No | - | 0.99% |
| 2 | CA2 | Yes | Yes | Yes | Yes | No | - | 0.55% |
| 3 | CA3 | No | - | No | - | No | - | - |
| 4 | CA4 | No | No | Yes | Yes | No | - | 0.42% |
| 5 | CA5 | Yes | Yes | No | Yes | No | - | 0.17% |
| 6 | CA6 | Yes | Yes | Yes | Yes | No | - | 0.46% |
| 7 | CA7 | No | - | No | - | No | - | - |
| 8 | CA8 | No | - | No | - | No | - | - |
| 9 | CA9 | No | - | No | - | No | - | - |
| 10 | CA10 | No | - | No | - | No | - | - |
| 11 | CA11 | No | i.s. | No | i.s. | Yes | i.s. | - |
| 12 | CA12 | No | i.s. | No | i.s. | Yes | No | - |
| 13 | CA13 | No | i.s. | Yes | i.s. | No | i.s. | - |
| 14 | CA14 | No | i.s. | Yes | i.s. | No | i.s. | - |
| 15 | CA15 | Yes | Yes | No | No | Yes | No | 0.33% |
| 16 | CA16 | No | Yes | Yes | Yes | No | No | 0.21% |
| 17 | CA17 | Yes | Yes | Yes | Yes | Yes | Yes | 0.77% |
| 18 | CA18 | Yes | Yes | Yes | Yes | Yes | Yes | 1.28% |
| 19 | CA19 | Yes | Yes | No | Yes | Yes | Yes | 0.67% |
| 20 | CA20 | Yes | Yes | No | Yes | No | Yes | 0.33% |
| 21 | SA1 | Yes | Yes | No | No | No | - | 0.52% |
| 22 | SA2 | No | No | Yes | Yes | No | - | 0.19% |
| 21 | SCH1 | Yes | Yes | No | No | Yes | No | 14.90% |
| 22 | SCH2 | Yes | Yes | Yes | No | No | - | 25.30% |
| 23 | LCH | No | No | No | No | No | No | - |
| 24 | AVM | No | No | No | No | No | No | - |
| 25 | VM | No | No | No | No | Yes | No | - |

### Validation of IDH1/2 mutant endothelial cells

To confirm the presence of the mutations, we performed BDA-PCR. After confirming that BDA-PCR promoted the preferential amplification of an *IDH1* hotspot mutations from SCH (Fig. 1A), we determined that 55% of angiosarcoma have *IDH1/2* hotspot mutations by both NGS and BDA-PCR, while none of the other vascular tumors, including 8 additional vascular neoplasms assessed by BDA-PCR alone, possessed any such mutations (Extended Data Table 1). Interestingly, BDA-PCR revealed additional *IDH1/2* hotspot mutations in several angiosarcoma cases, which were not identified by amplicon NGS, consistent with the higher reported sensitivity of BDA-PCR compared with amplicon NGS (Table 1; Extended Data Table 1).(23)

**Figure 1.**
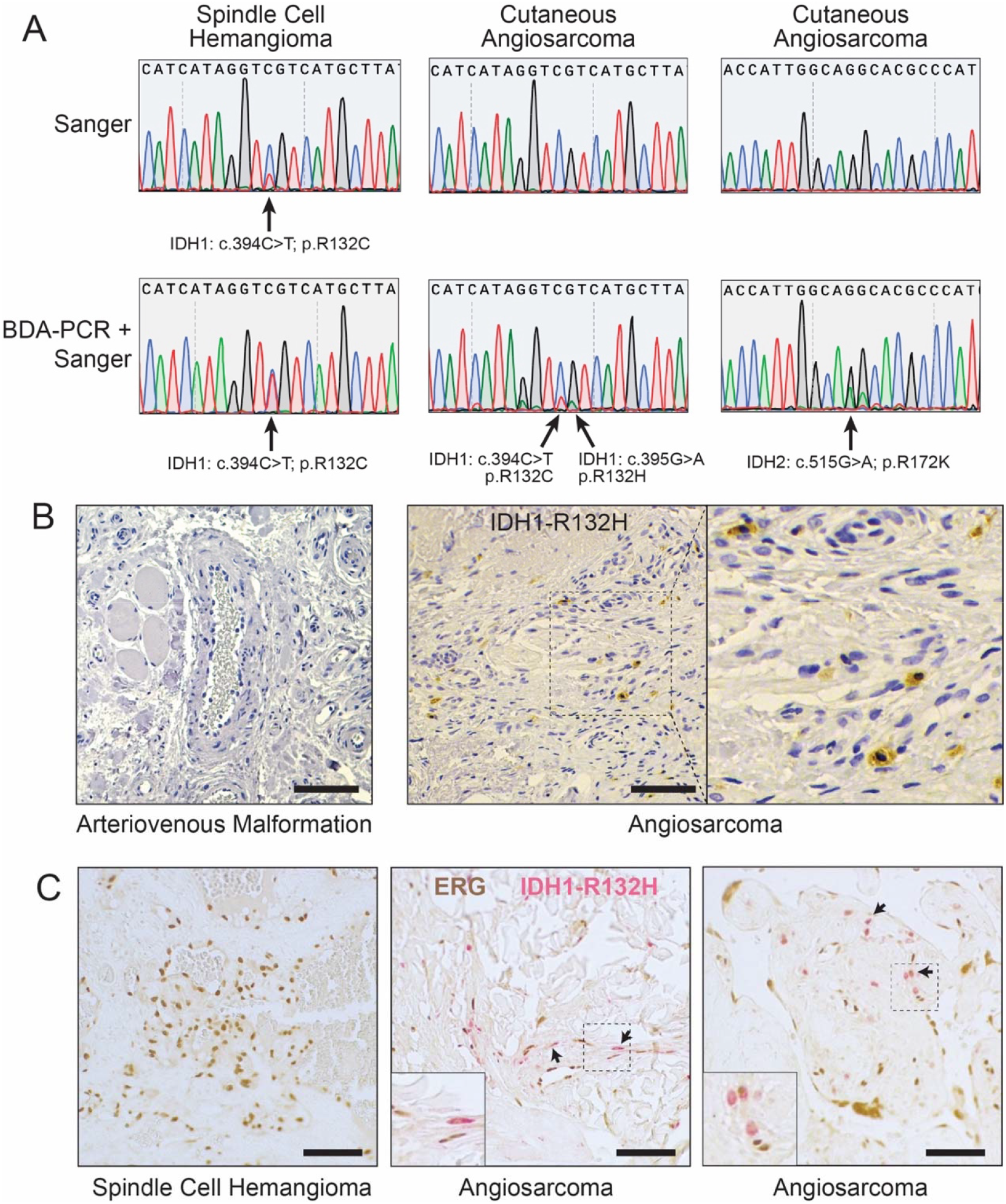
Validation of subclonal *IDH1/2* mutations in vascular tumors. (A) Blocker displacement amplification PCR (BDA-PCR) products showing enrichment of mutant alleles. Middle panels (case CA18) confirms the presence of both *IDH1* R132C and R132H mutant alleles in the same angiosarcoma sample. (B) Immunohistochemistry (IHC) confirms strong cytoplasmic and weak nuclear expression of IDH1 R132H in a subset of cells in cutaneous angiosarcoma (left, negative control, arteriovenous malformation (AVM); middle and right, CA2). (C) Dual IHC reveals concomitant nuclear ERG (brown) and cytoplasmic IDH R132H (magenta) expression in cutaneous angiosarcoma (left, negative control, SCH1; middle, CA14; right, CA1). Bar, 100 μm.

An antibody specific for IDH1 R132H was used to confirm expression of mutant IDH1 in a subset of samples with R132H mutations by sequencing analysis. IHC confirmed strong cytoplasmic and weak nuclear expression of IDH1 R132H in a fraction of cells in angiosarcoma samples, but not in SCH or other vascular tumors (Fig. 1B; Extended Data Fig. 3). Given the frequent infiltration of angiosarcoma into normal tissue, we sought to confirm that the IDH mutations were occurring in angiosarcoma cells, rather than interstitial cells or leukocytes. Dual staining for ERG, an endothelial cell marker, revealed cells that stained positively for both ERG in the nucleus and IDH1 R132H in the cytoplasm, demonstrating that IDH1 R132H expression could be detected in a subset of the malignant endothelial cells in angiosarcoma (Fig. 1C). Thus, through amplicon sequencing, BDA-PCR, and IHC, we identified the presence of *IDH1/2* mutations in angiosarcoma.

### Non-cell-autonomous effects of IDH1 mutations

Clinical studies of *IDH1/2*-mutant inhibitors in acute myeloid leukemia and cholangiocarcinoma have shown that inhibition of these enzymes can provide therapeutic benefit, even when mutations are present at low VAF.(24-26) To test whether subclonal mutations in *IDH1/2* might have a functional role in angiosarcoma, *IDH1* R132C and R132H mutants were cloned and transduced into hTERT-immortalized human aortic endothelial cells (TeloHAEC), and expression of the mutant alleles was confirmed by Western blot (Extended Data Fig. 4A). Expression of IDH1 mutants did not significantly impact cell proliferation (Extended Data Fig. 4B). Given the low VAF of *IDH1/2* mutation in the angiosarcomas, we hypothesized that the mutations might exert non-cell autonomous effects on endothelial cells, likely through the secretion of D-2-HG. *IDH1/2* mutations have been reported to affect hypoxia inducible factor (HIF) signaling,(27-29) so we tested its non-cell autonomous effects on HIF signaling in endothelial cells. Pilot experiments using TeloHAECs treated with D-2-HG revealed a significant induction of *VEGF* and *SLC2A1 (GLUT1)* expression and robust activation of Akt and ERK signaling by the oncometabolite (Extended Data Fig. 5A,B). Conditioned media was harvested from TeloHAECs transduced with *IDH1* R132C and R132H, or from *IDH1* R132C-expressing HT1080 cells (Extended Data Fig. 4C,D). As expected, conditioned media from *IDH1* R132C and R132H-tranducedTeloHAECs showed elevated levels of 2-HG. Ivosidenib, a small molecule inhibitor that specifically binds and inhibits mutant IDH1, inhibited 2-HG accumulation (Fig. 2A). After incubation of parental TeloHAEC cells with conditioned medium, the recipient TeloHAECs showed significantly increased expression of *VEGF* and *SLC2A1* (*GLUT1*) (Fig. 2B; Extended Data Fig. 4E). TeloHAECs treated with conditioned media also showed robust activation of pAkt and pERK under growth-factor free conditions (Extended Data Fig. 4C; Extended Data Fig. 5C). Notably, conditioned media derived from ivosidenib-treated donor cells rescued both transcriptional changes and activation of Akt and ERK, implicating D-2-HG as a likely effector driving the non-cell autonomous effects of *IDH1* mutations in endothelial cells (Fig. 2A-C). Since *IDH1/2* mutations have also been reported to increase DNA methylation(30), we next tested whether conditioned media from *IDH1* R132C expressing cells could induce DNA methylation changes in recipient cells. Treatment with R132C-conditioned media induced notable changes in CpG methylation, which was increased overall when compared to parental cells. As expected, this increase was rescued by pre-treatment of R132C donor cells with ivosidenib (Fig. 2D).

**Figure 2.**
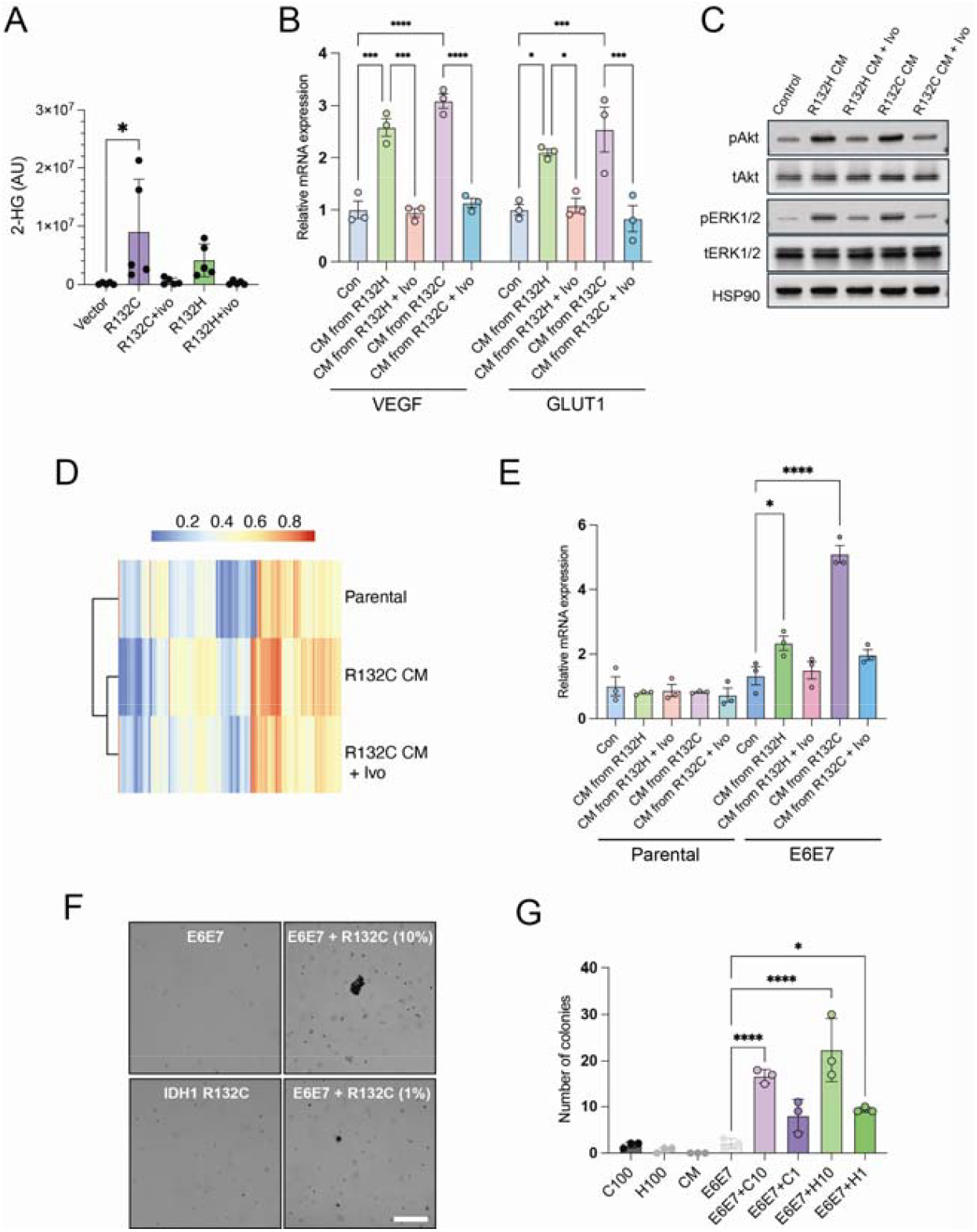
*IDH1* mutant alleles can exert non-cell autonomous effects on endothelial cells. (A) Relative levels of 2-HG in media of indicated cell type as measured by LC-MS/MS (*n*=5). (B) Transcriptional induction of pro-angiogenic (*VEGF*) and metabolic (*SLC2A1/GLUT1*) genes in wild-type cells by conditioned media (CM) from mutant cells, reversed by 2 μM ivosidenib (Ivo) pretreatment (*n*= 3). (B) Western blot analysis of TeloHAECs incubated with CM from R132H or R132C mutant cells for 48 hours followed by 2 h incubation in growth-factor free media. Activation of pAkt and pERK1/2 induced by CM is inhibited by pretreatment of donor cells with ivosidenib. (D) Heatmap depicting unsupervised clustering of CpG sites showing greatest variance in TeloHAECs treated with IDH1 R132C CM. The beta value for each probe ranges from 0 (blue, unmethylated) to 1 (red, fully methylated). (E) Relative mRNA expression of ACTA2 (α-SMA) following CM treatment ± 2 μM ivosidenib pretreatment of donor cells. (F) Representative images of soft agar colony formation assays performed using E6E7 TeloHAECs alone or co-cultured with IDH1 mutant cells at defined ratios. Bar, 500 μm. (G) Quantification of colony numbers from soft agar assays under the indicated cell mixture conditions. Bar graphs show mean ± s.e.m. from three independent biological replicates (n = 3). Statistical analysis was performed using two-way ANOVA with Šídák’s multiple comparisons test (B, E) or one-way ANOVA with multiple comparisons (A. G). Exact P values are provided in the Source Data. *P < 0.05, ***P < 0.001, ****P < 0.0001.

Consistent with multistage endothelial transformation models, E6/E7-immortalized TeloHAECs were used as a permissive background to test whether non-cell-autonomous signals from IDH1-mutant cells might provide additional oncogenic signals required to transform endothelial cells.(31,32) Conditioned media (CM) from IDH1 R132H- or R132C-expressing donor cells increased *VEGF* and *SLC2A1* (GLUT1) expression in recipient endothelial cells and activated AKT and ERK signaling, as assessed by pAKT and pERK1/2 immunoblotting (Extended Data Fig. 6A,B). Pretreatment of donor cells with ivosidenib during CM generation rescued these transcriptional and signaling responses (Extended Data Fig. 6A,B). In the same CM framework, IDH1-mutant CM induced an EndMT-associated transcriptional program— including ACTA2 (α-SMA), ZEB1, SNAI1, S100A4/FSP1 and VIM—that was largely absent in parental TeloHAECs but robustly induced in E6/E7-immortalized cells (Fig. 2E; Extended Data Fig. 7). Next, we tested the impact of mutant IDH1 signaling on anchorage-independent growth. Soft-agar assays were performed using E6/E7 TeloHAECs cultured alone or co-cultured with IDH1-mutant cells at defined mixture ratios (1% or 10% mutant cells, with the remainder comprised of E6/E7 cells). Under these conditions, co-culture with IDH1-mutant cells significantly increased colony formation relative to any individual cell type alone after 4 weeks (Fig. 2F-G, Extended Data Fig. 8). Together, these findings indicate that low-VAF IDH1 mutations can exert non-cell-autonomous, transforming effects on endothelial cells. In a pre-neoplastic endothelial cell context, these effects extend to EndMT-associated transcriptional reprogramming and development of anchorage-independent colony formation during direct co-culture.

### Clinical response to ivosidenib

Given ivosidenib’s safety profile and its efficacy in treating chondrosarcoma with subclonal IDH1 mutations,(33) ivosidenib was administered to three patients with mosaic IDH1 R132C mutations (Extended Data Table 2). Patient 1 was a 35-year-old female with progressive SCH and enchondromas for 25 years, consistent with Maffucci syndrome, refractory to surgery, compression and sirolimus. Ivosidenib (500 mg daily) led to reduced SCH size with decreased pain and improved mobility by ∼3 months (Fig. 3A), discontinuation of scheduled pain medications by 6 months, and reduced tumor hardness by SkinFibrometer (Fig. 3B). Serum D-2-HG decreased from 362 ng/mL to 96 ng/mL (normal) within 1 week and remained normal (Fig. 3C), without significant drug-related adverse events over 18 months. Patient 2 was a 34-year-old female who developed painful SCH starting at age 7, with recurrence after excisions and sirolimus. After initiating ivosidenib (500 mg daily), she experienced rapid improvement in pain and mobility (Fig. 3D) and remained on therapy for 12 months with continued improvement and no persistent side effects requiring dose adjustment. Patient 3 had congenital lymphedema and was diagnosed with angiosarcoma at age 19 after biopsy of a violaceous right lower leg lesion, treated with below-knee amputation and adjuvant chemotherapy (doxorubicin, ifosfamide, paclitaxel). At age 23, he developed recurrent Dabska tumors managed with resection, sequencing revealed IDH1 R132C with negative germline testing, consistent with mosaicism. He later developed painful progressive osseous lesions (T1, T9, right 9th rib; Fig. 3E) that harbored IDH1 R132C but showed atypical vascular lesions without angiosarcoma on biopsy. After no improvement on a broad-spectrum tyrosine kinase inhibitor, ivosidenib (500 mg daily) resulted in pain resolution and increased sclerosis consistent with response (Fig. 3E), with durable control for >2 years and only mild intermittent grade 1 diarrhea.

**Figure 3.**
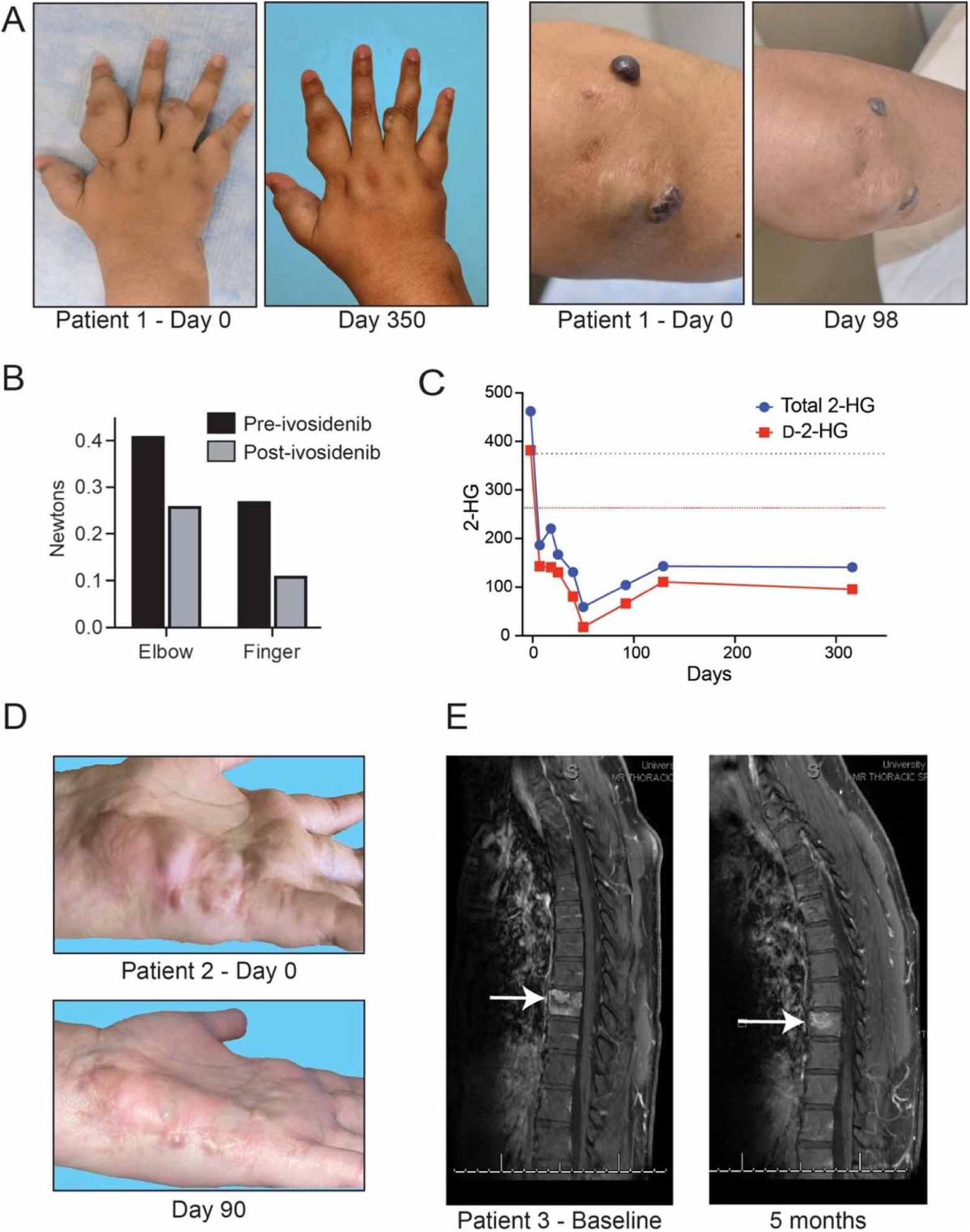
Clinical response to ivosidenib in patients with mosaic *IDH1* mutation (A) Images of SCH on elbow and forearm after 0 days and 98 days of treatment with ivosidenib (SCH1). (B) Tumor induration decreased 2 months after initiation of ivosidenib (Delfin SkinFibrometer). (C) Decrease in serum concentration of total 2-HG and D-2-HG that was maintained for the duration of treatment. Initial serum 2-HG measurement on Day -2. Horizontal lines indicate upper limit of normal for total 2-HG (blue) and D-2-HG (red). (D) Hemangiomas on the hands after 0 days and 90 days of treatment with ivosidenib (SCH2). (E) Ivosidenib induces sclerosis and decreases the size of an atypical vascular lesion with *IDH1* R132C mutation.

## Discussion

We describe the discovery of low VAF, hotspot mutations in *IDH1* and *IDH2* in 55% of angiosarcomas. The combination of amplicon sequencing, which had a median read depth of >5000x for *IDH1/2* and could detect VAF at 0.16%, and BDA-PCR, which could detect VAF at <0.1%,(34) allowed us detect *IDH1/2* mutants, despite their low VAF (0.17-1.28%). Only mutations detected by amplicon sequencing and validated by BDA-PCR/Sanger were considered *IDH1/2* mutant, so our study may have underestimated the true prevalence of *IDH1/2* mutations in angiosarcoma. The low VAF of the mutations detected here, differences in assays used, and the infiltrative properties of angiosarcoma may explain why *IDH1/2* mutations have not been detected in prior sequencing studies.(2,9,20,21) Strikingly, more than one-quarter of the angiosarcomas in this study possessed multiple distinct *IDH1/2* hotspot mutations within the same tumor; in several cases, three different gain-of-function mutations were detected. The high prevalence of *IDH1/2* mutations in angiosarcoma suggests that these mutations may play a role in the development of this cancer. Moreover, the availability of mutant IDH inhibitors raises the possibility that *IDH1/2* mutations in angiosarcoma may have a therapeutic role.

Specific oncogenic mutations (e.g. *KRAS, PIK3CA* activating mutations) can exert biological effects, even when they are present at low VAF, especially in vascular lesions.(35-37) Moreover, *IDH1/2* has already been reported to be oncogenic at subclonal frequencies in a other cancers.(23,26,38-40) We provide evidence that low VAF *IDH1/2* mutations may exert non-cell-autonomous transforming effects through the secretion of D-2-HG. Consistent with studies in astrocytes(28,41), *IDH1/*2 mutation did not provide endothelial cells with a cell autonomous growth advantage in vitro. However, conditioned media from IDH1-mutant expressing cells induced VEGF expression, activated Akt and Erk signaling, and increased DNA methylation, all of which were rescued by pre-treatment of *IDH1/2* mutant cells with ivosidenib. Our study contributes further evidence that D-2-HG promotes transformation as a diffusible “oncometabolite” through its pleiotropic effects on glucose sensing and metabolism, HIF-1α signaling, DNA and histone methylation, DNA damage, and immunomodulation.(19,28,42,43) The E6/E7-immortalized endothelial system provides a neoplastic precursor to test the impact of non-cell-autonomous signaling readouts on transformation-associated phenotypes. While CM elicited shared paracrine responses across endothelial settings (for example, VEGF/GLUT1 induction and Akt/Erk activation), EndMT-associated transcriptional changes were mostly noted in the E6/E7-immortalized setting, enabling functional testing of cooperative interactions on anchorage-independent growth. Our findings are consistent with animal models in which minority subpopulations within genetically heterogenous tumors can play critical roles in tumorigenesis through paracrine signaling.(6,10,44-47) In an endothelial context, our co-culture experiments provide functional support for this concept. The detection of *IDH1/2* mutations in most mature tumors suggests that the persistent ability to secrete D-2-HG may be required not only to promote tumor development but also for angiosarcoma persistence.

This model is bolstered by the observation that both sporadic and Maffucci-associated SCH arise from subclonal *IDH1/2* mutations, with VAFs ranging from ∼4-25% (Table 1).(4,48) Extending previously reported cases of the successful treatment of subclonal-*IDH1* mutant tumors and enchondromas with ivosidenib, we report striking clinical responses to ivosidenib in patients with mosaic *IDH1* mutations who presented with atypical vascular lesions, including patients with Maffucci syndrome.(12) While the patients described here showed striking clinical improvements and minimal side effects, clinical trials and long-term follow-up are required to evaluate the safety and long-term efficacy of ivosidenib in treating Maffucci syndrome. Additional in vivo studies are also necessary to dissect the specific roles of low frequency *IDH1/2* mutations in the pathogenesis of angiosarcoma, and potentially other soft tissue sarcomas. If *IDH1/2* mutations are found to be required for the progression of angiosarcomas, the addition of mutant *IDH1/2* inhibitors to current standard-of-care therapies should be tested, as they may offer benefits for patients with advanced disease.

## Methods

### Study Design

This study was approved by the University of Texas Southwestern (STU 072018-067) and University of Colorado Institutional Review Boards (COMIRB 25-1485 and 15-1461). Archived, formalin-fixed, paraffin-embedded (FFPE) tissues from patients with cutaneous angiosarcoma, splenic angiosarcoma, and control vascular tumors collected between January 1, 2015, and May 1, 2025, were utilized. Because specimens were obtained during standard-of-care treatment, the study was deemed exempt from the requirement for written informed consent.

### Targeted Amplicon Sequencing and Data Analysis

FFPE slides were submitted for DNA extraction and targeted amplicon sequencing using the Paragon CleanPlex OncoZoom Cancer Hotspot panel (Admera Health). Sequencing reads were quality- and adapter-trimmed using fastp and aligned to the hg38 reference genome using bowtie2 (very-sensitive-local). Median coverage across the IDH1 and IDH2 exons of interest was 5,599× (range, 3,839–9,955×). Low-frequency variants in IDH1 and IDH2 were detected using both Samtools (RRID:SCR_002105) and LoFreq (RRID:SCR_013054), whereas variants in other genomic regions were identified using LoFreq alone with default parameters. Variants called by Samtools were filtered based on a conservatively estimated PCR error rate derived from the number of amplification cycles (10–28 cycles; 0.0016–0.0021 of reads containing one mutation). Variant annotation was performed using vcf2maf2 and VEP. Variants outside IDH1 and IDH2 occurring at >1% allele frequency and present at <0.05% in any global population database were also identified. Only mutations previously reported as cancer hotspots (>10 occurrences in COSMIC (RRID:SCR_002260) or cBioPortal) were reported in Extended Data Table 1. BLASTn (v2.12.0, RRID:SCR_001598) was used to align amplicon sequencing reads to mutant IDH1 R132C, IDH1 R132H, or IDH2 R172K reference sequences (Extended Data Table 3). To exclude allelic double mutations, a combined IDH1 R132C/R132H reference sequence was also tested. Reads aligning for at least 25 consecutive bases across the single-nucleotide polymorphism without mismatches were retained and quantified. Mutant reads were compared to wild-type IDH1 or IDH2 reference sequences, and random base substitutions were used as negative controls to assess background signal.

### Blocker Displacement Amplification (BDA)-Polymerase Chain Reaction

Blocker displacement amplification polymerase chain reaction (BDA-PCR) was performed as previously described with minor modifications(23,34). A Taq-based, non–error-correcting polymerase (Sapphire Master Mix, Takara) was used for both standard PCR and wild-type blocking reactions. Standard PCR was performed using 20 ng of DNA with 0.1 µM forward and reverse primers, with cycling conditions of 95°C for 3 min followed by 30 cycles of 95°C for 10 s and 60°C for 2 min. For BDA-PCR, a primer mix containing forward, reverse, and blocker oligonucleotides was prepared as described. For each reaction, 2.5 µL of primer mix was combined with 100 ng of DNA in a 25 µL reaction. Cycling conditions consisted of 95°C for 3 min; 20 cycles of 95°C for 30 s, 68°C for 15 s with a 0.5°C decrease per cycle, and 72°C for 1 min; followed by 16 cycles of 95°C for 30 s, 58°C for 15 s, and 72°C for 1 min, with a final extension at 72°C for 10 min. PCR products were purified and submitted for Sanger sequencing.

### Whole Genome Amplification

When DNA quantities for BDA-PCR were <100 ng, samples were first amplified using the GenomePlex Whole Genome Amplification kit (Sigma, WGA1). PCR products were purified using the NucleoSpin PCR & Gel Clean-Up kit (Takara, 740609.250). For fragmentation and library preparation, 10 µL of DNA at 1 ng/µL was used as recommended. When initial DNA concentration was <1□ng/µL, 10 ng of DNA was added in a larger volume and other reaction components were scaled proportionally. Briefly, genomic DNA was mixed with 10× Fragmentation Buffer, incubated at 95°C for exactly 4□min, and immediately cooled on ice. Library preparation was performed by adding 1× Library Preparation Buffer and Library Stabilization Solution, incubating at 95°C for 2□min, then cooling on ice, followed by addition of Library Preparation Enzyme. The reaction was incubated sequentially at 16°C for 20□min, 24°C for 20□min, 37°C for 20□min, and 75°C for 5□min. Library amplification was subsequently carried out using JumpStart Taq DNA Polymerase (Sigma, D9307, RRID:SCR_000488) with a thermocycler profile of 95°C for 3□min, followed by 15 cycles of 95°C for 15□s and 65°C for 5□min. PCR products were purified using the NucleoSpin PCR & Gel Clean-Up kit (Macherey-Nagel, Düren, Germany), either directly or after separation on a 1.8% agarose gel and gel excision prior to BDA-PCR.

### Immunohistochemistry

FFPE sections were deparaffinized, heat retrieved (Citrate buffer, Epredia, AP-9003-125, 95°C for 30 min), permeabilized (0.2% Triton X-100 for 10 min), and blocked in 3% BSA. For single-staining of IDH1 R132H, these sections were then incubated overnight at 4°C with prediluted IDH1 R132H primary antibody (Sigma, 456R-37). On the following day, slides were stained with the appropriate secondary antibody (Invitrogen, G-21234) and developed with the DAB substrate kit (Vector, SK-4100). After confirmation of the antibody, automated IHC staining was completed using a Dako stainer. For dual IHC, both IDH1 R132H and ERG () were applied to the same section, with ERG detected using its primary antibody (Biocare Medical) and a mouse secondary antibody (ThermoFisher, 31430), and chromogenic visualization performed with the Vector dual-stain kit (MP-7724-15). Nuclei were visualized by counterstaining with hematoxylin (Vector, H-3401-500).

### *IDH1* Mutant Cloning and Expression

*IDH1* wild type, *IDH1* R132C, and *IDH1* R132H cDNA were synthesized as gBlock gene fragments (Integrated DNA Technologies), PCR amplified using Q5 high-fidelity polymerase (NEB, M0492S) and cloned into pLenti-6.3 MCS vector using BamHI and SalI restriction digest. All constructs were confirmed by Sanger or whole plasmid sequencing.

### Cell Culture and Lentiviral Transduction

Telomerase immortalized human aortic endothelial cells (TeloHAEC) were cultured in Endothelial Cell Media supplemented with 2% dialyzed FBS (Corning, MT35071CV) and 1× Antibiotic-Antimycotic solution. Conditioned media (CM) was generated from donor TeloHAECs expressing wild-type or mutant IDH1 or HT1080 cells (RRID:CVCL_0317). Donor cells were seeded one day prior to treatment, and the culture medium was replaced the next day with fresh growth media containing either vehicle or ivosidenib (Ivo). After 72 h, CM was collected and clarified by centrifugation at 1,500 g for 15 min, followed by filtration through a 0.22-µm membrane. Recipient TeloHAECs were plated the day before treatment and incubated with donor-derived CM for 48 hours. Following CM exposure, cells were harvested for downstream assays, including immunoblotting and quantitative RT–PCR. For D-2-hydroxyglutarate (D-2-HG) treatments, TeloHAECs were incubated with D-2-HG (800 μM) for 48 hours. At the end of treatment, cells were washed and the medium was replaced with growth factor– and serum-free endothelial media. Whole-cell lysates were collected at 0, 15, 60, and 120 min after media replacement to assess time-dependent activation of downstream signaling pathways.

For generating lentiviruses, LentiX-293T cells (Clontech, 632180) were seeded at approximately 60% confluence in antibiotic-free media 12 to 16 hours before transfection. shRNA (4.5 μg) or expression plasmid (4.5 μg), 2.5 μg of pMD2.G (RRID:Addgene_12259), and 4.5 μg of psPAX2 (RRID:Addgene_12260) were co-transfected into LentiX-293T cells using Lipofectamine 3000 (Thermo Fisher Scientific, L3000015) according to the manufacturer’s protocol. Viruses were collected after 48 h after transfection. For lentiviral transduction, cells were seeded into 10-cm dishes at 70% confluence. Viral supernatant was added to the cells in the presence of polybrene (8 μg/mL). Following transduction, cells were selected with Blasticidin at 5 μg/mL.

### Immunoblotting

For immunoblot analyses, cells were lysed in Cell Lysis Buffer (Cell Signaling Technology, 9803) with Protease/Phosphatase inhibitor (Cell Signaling Technology, 5872), and whole-cell lysates were mixed with Laemmli Sample Buffer (Bio-Rad, 1610747) prior to SDS–PAGE. Proteins were separated by electrophoresis, transferred onto nitrocellulose membranes, and probed with primary antibodies against IDH1 (Proteintech, 12332-1-AP), IDH1 R132H (Sigma, 456R-37), p-AKT (Cell Signaling Technology, 4060), p-ERK (Cell Signaling Technology, 4695), total AKT (Cell Signaling Technology, 2920), total ERK (Cell Signaling Technology, 4370), and Hsp90 (Cell Signaling Technology, 4877). After incubation with HRP-conjugated secondary antibodies, proteins were visualized using Western Lightning Plus-ECL (PerkinElmer, 50-904-9323).

### RNA Extraction and qRT–PCR

Total RNA was isolated from cultured cells using the RNeasy Mini Kit (Qiagen, 74106). Extracted RNA was reverse transcribed into cDNA using the iScript cDNA Synthesis Kit (Bio-Rad, 1708891) following the manufacturer’s instructions. Quantitative RT–PCR was performed using PowerUp SYBR Green Master Mix (Applied Biosystems, A25779) with gene-specific primers (Extended Data Table 2).

### DNA Methylation Analysis

Genomic DNA (>5 ug) was isolated using a Qiagen DNA extraction kit (Qiagen, 51304) according to the manufacturer’s instructions. Bisulfite treatment was performed using the EZ DNA Methylation kit (Zymo Research), which included internal positive and negative control samples. The bisulfite-converted samples were processed using the Illumina Infinium Methylation assay using the Infinium Epic-8 v2.0 BeadChips, which includes ∼850,000 single CpG sites (Illumina Inc, San Diego, CA). The labeling, hybridization, and scanning procedures were performed on the Illumina IScan system in the University of Colorado Genomics Core. DNA methylation array data was processed using the minfi R package (v1.56.0). Probes with insufficient signal, as defined by a detection p-value < 0.01, were removed from the data set and the data was normalized using the Funnorm method. The heatmap was generated using the top 2,000 genes with the highest variance between the parental and IDH R132C mutant and dendrograms were generated using hierarchical clustering.

### Colony formation (soft agar) assay

Anchorage-independent growth was assessed by soft-agar colony formation. Base layers (0.5% agar) were prepared in DMEM (diluted from 2× DMEM) supplemented with 12% fetal bovine serum (FBS) and penicillin/streptomycin/amphotericin (P/S/A). Top layers (0.3% agar) containing cells were overlaid onto the base layer. A total of 15,000 cells were plated per well in triplicate. For co-culture conditions, IDH1-mutant cells were mixed with E6/E7 TeloHAECs at the indicated percentages (e.g., C1 and C10 denote 1% or 10% R132C cells, respectively; H1 and H10 denote 1% or 10% R132H cells), with the remainder comprised of E6/E7 cells. Cultures were maintained for 4 weeks, with media changes twice weekly (1 mL per well) using DMEM supplemented with 10% FBS. For the indicated condition, media changes were performed twice weekly using 800 µL R132C conditioned media plus 200 µL DMEM with 10% FBS.

### Ivosidenib Treatment

Written informed consent was obtained from the patients. Ivosidenib was provided by Servier through the patient assistance program. Patients received 500 mg of oral ivosidenib daily. Assessments included photographs, assessment of hemangioma pain and size, measurements of tumor induration as taken by SkinFibrometer (Delfin), serum D-2-HG and total 2-HG levels, and CT and x-rays of enchondromas when relevant. Background images in some photos were replaced in Adobe Photoshop (RRID:SCR_014199). Measurements were repeated every 30 days for 4 months following initiation of ivosidenib treatment. To evaluate for safety and adverse events, an ECG to monitor for QT prolongation was performed and CK, LD, CMP, and CBC labs were drawn weekly for one month and monthly thereafter.

## Data availability

DNA methylation array and targeted amplicon sequencing data were generated in this study. These data will be available upon publication. Access to clinical data may be considered upon reasonable request to the corresponding author and subject to institutional review board approval. All other data supporting the findings of this study are included within the paper or its Extended Data.

## Code availability

Code used to process, quality control, and generate figures for the DNA methylation analysis is available in a public repository: https://github.com/KatyW15/Angiosarcoma_DNA_methylation_analysis.

## Acknowledgements

This project was supported by the National Cancer Institute (5R01CA275071 to RCW; ZIA BC 011894 to GJS), the University of Colorado Cancer Center Genomics/Microarray Core (P30CA046934), research funding from Servier BioInnovation (RCW), and startup funds from the University of Colorado Anschutz Medical Campus (RCW). The contributions of the NIH author are considered works of the United States Government. The findings and conclusions presented in this paper are those of the author and do not necessarily reflect the views of the NIH or the U.S. Department of Health and Human Services. We thank the participants with mosaic IDH1 mutation for agreeing to participate in this study. We thank Dr. Lu Le for use of the SkinFibrometer; Dr. Phil Shaul for the gift of TeloHAEC cells; and Dr. David Schwartz for comments on the manuscript.

