## Supplementary Data for "Subclonal IDH1/2 Mutations as a Targetable Vulnerability in Vascular Tumors"

| **Extended Data Table 1. Detailed summary of *IDH1* and *IDH2* mutations in angiosarcoma and other vascular tumors** | | | | | | | | | | | | | | | | | | |
| --- | --- | --- | --- | --- | --- | --- | --- | --- | --- | --- | --- | --- | --- | --- | --- | --- | --- | --- |
| **Case** | **Diagnosis** | **Age/**  **Gender** | **Location** | **IDH1 R132C** | | | | **IDH1 R132H** | | | | **IDH2 R172K** | | | |  | ***IDH1/2***  **VAF (%)** | **Additional Mutations;**  **Tumor Cellularity**  **(low, moderate, high)** |
|  |  |  |  | **NGS** | **BDA-PCR** | **Lofreq** | **BLAST** | **NGS** | **BDA-PCR** | **Lofreq** | **BLAST** | **NGS** | **BDA-PCR** | **Lofreq** | **BLAST** |  |  |  |
| **1** | CA1 | 67/M | Scalp | Yes  (0.64) | Yes | Yes  (0.63) | Yes | Yes  (0.35) | Yes | Yes  (0.35) | Yes | No | - | Yes  (0.15) | Yes |  | 0.99 | moderate |
| **2** | CA2 | 82/F | Nose | Yes  (0.24) | Yes | No | Yes | Yes  (0.31) | Yes | Yes  (0.31) | Yes | No | - | No | No |  | 0.55 | high |
| **3** | CA3 | 80/M | Neck | No | - | No | No | No | - | No | No | No | - | No | No |  | - | high |
| **4** | CA4 | 83/M | Scalp | No | No | No | No | Yes  (0.42) | Yes | Yes  (0.42) | Yes | No | - | No | No |  | 0.42 | high |
| **5** | CA5 | 83/F | Scalp | Yes  (0.17) | Yes | No | Yes | No | Yes | No | Yes | No | - | No | - |  | 0.17 | high |
| **6** | CA6 | 85/F | Nose | Yes  (0.23) | Yes | No | Yes | Yes  (0.23) | Yes | No | Yes | No | - | No | No |  | 0.46 | moderate |
| **7** | CA7 | 89/F | Eyebrow | No | - | No | No | No | - | No | No | No | - | No | No |  | - | *TP53* X187splice;  PIK3CA p.H1047R; MAP2K1 p.E203K;  moderate |
| **8** | CA8 | 82/M | Scalp | No | - | No | No | No | - | No | No | No | - | No | No |  | - | moderate |
| **9** | CA9 | 80/M | Cheek | No | - | No | No | No | - | No | No | No | - | No | No |  | - | moderate |
| **10** | CA10 | 80/M | Scalp | No | - | No | No | No | - | No | No | No | - | No | No |  | - | low |
| **11** | CA11 | 69/M | Forehead | No | i.s. | No | No | No | i.s. | No | No | Yes | i.s. | Yes  (0.16) | Yes |  | - | moderate |
| **12** | CA12 | 79/F | Forehead | No | i.s. | No | No | No | i.s. | No | No | Yes  (0.73) | No | Yes  (0.72) | Yes |  | - | low |
| **13** | CA13 | 42/F | Scalp | No | i.s. | No | Yes | Yes | i.s. | No | No | No | i.s. | No | No |  | - | moderate |
| **14** | CA14 | 71/M | Forehead | No | i.s. | No | No | Yes  (0.42) | i.s. | Yes  (0.41) | Yes | No | i.s. | No | No |  | - | low |
| **15** | CA15 | 90/F | Cheek | Yes  (0.33) | Yes | No | Yes | No | No | No | No | Yes  (0.21) | No | Yes  (0.21) | No |  | 0.33 | low |
| **16** | CA16 | 100/M | Scalp | No | Yes | No | No | Yes  (0.21) | Yes | No | No | No | No | No | No |  | 0.21 | moderate |
| **17** | CA17 | 81/F | Cheek | Yes  (0.23) | Yes | No | Yes | Yes  (0.24) | Yes | No | Yes | Yes  (0.26) | Yes | Yes  (0.26) | Yes |  | 0.77 | low |
| **18** | CA18 | 86/M | Eyebrow | Yes  (0.60) | Yes | Yes  (0.60) | Yes | Yes  (0.40) | Yes | Yes  (0.40) | Yes | Yes  (0.28) | Yes | Yes  (0.27) | Yes |  | 1.28 | high |
| **19** | CA19 | 85/M | Scalp | Yes  (0.41) | Yes | No | Yes | No | Yes | No | Yes | Yes  (0.26) | Yes | Yes  (0.26) | Yes |  | 0.67 | HRAS p.Q61L;  high |
| **20** | CA20 | 81/M | Scalp | Yes  (0.33) | Yes | No | Yes | No | Yes | No | Yes | No | Yes | No | Yes |  | 0.33 | high |
| **21** | SA1 | 80/F | Spleen | Yes  (0.52) | Yes | Yes  (0.52) | Yes | No | No | No | No | No | - | No | No |  | 0.52 | moderate |
| **22** | SA2 | 48/F | Spleen | No | No | No | No | Yes  (0.19) | Yes | No | Yes | No | - | No | No |  | 0.19 | high |
| **21** | SCH1 | 35/F | Hand | Yes  (14.90) | Yes | Yes  (14.90) | Yes | No | No | No | No | Yes | No | Yes  (0.38) | Yes |  | 14.90 | KRAS p.D33N |
| **22** | SCH2 | 34/F | Hand | Yes  (25.30) | Yes | Yes  (25.30) | Yes | Yes | No | No | No | No | - | No | No |  | 25.30 |  |
| **23** | LCH | 44/F | Chest | No | No | No | No | No | No | No | No | No | No | No | No |  | - |  |
| **24** | AVM | 38/M | Lip | No | No | No | No | No | No | No | No | No | No | No | No |  | - | MAP2K1 p.K57N |
| **25** | VM | 65/M | Lip | No | No | No | Yes | No | No | No | Yes | Yes  (0.24) | No | Yes  (0.24) | Yes |  | - | PIK3CA p.H1047R |
| **26** | CHA1 | 43/F | Thigh | - | No | - | - | - | No | - | - | - | i.s. | - | - |  | - | - |
| **27** | CHA2a | 47/F | Lower chest | - | No | - | - | - | No | - | - | - | No | - | - |  | - | - |
|  | CHA2b |  | Lower abdomen | - | No | - | - | - | No | - | - | - | No | - | - |  | - | - |
| **28** | KS1a | 95/F | Medial foot | - | No | - | - | - | No | - | - | - | No | - | - |  | - | - |
|  | KS1b |  | Dorsal hand | - | No | - | - | - | No | - | - | - | No | - | - |  | - | - |
| **30** | VL1 | 72/F | Lip | - | No | - | - | - | No | - | - | - | No | - | - |  | - | - |
| **31** | VL2 | 69/F | Eyelid | - | No | - | - | - | No | - | - | - | i.s. | - | - |  | - | - |

‘-‘ BDA-PCR was not completed; ‘i.s.’ insufficient sample for BDA-PCR; ‘CA’ cutaneous angiosarcoma; ‘SA’ splenic angiosarcoma; ‘SCH’ spindle cell hemangioma; ‘LCH’ lobular capillary hemangioma; ‘AVM’ arteriovenous malformation; ‘VM’ venous malformation; ‘CHA’ cherry angioma; ‘KS’ Kaposi sarcoma; ‘LA’ lymphangioma; ‘VL’ venous lake; ‘Total VAF’ – Only mutant alleles that were confirmed by both NGS and BDA-PCR were included in the ‘total VAF’ sum.

| **Extended Data Table 2. Clinical characteristics and responses to ivosidenib in patients with mosaic IDH1 R132C mutations** | | | | | | |
| --- | --- | --- | --- | --- | --- | --- |
|  | Age/sex | Presentation | Prior therapies | Ivosidenib | Best response | Objective measures / Safety |
| 1 | 35/F | Worsening SCH + enchondromas (25 years); painful enlarging hemangiomas (hands) consistent with Maffucci syndrome | Surgery; compression; sirolimus (without success) | 500 mg daily; >20 months | ↓SCH size; ↓pain; ↑mobility (Fig. 3a); pain meds stopped by 6 months; continued improvement | Serum D-2HG 362→96 ng/mL at 1 week; sustained normal (Fig. 3c); SkinFibrometer: ↓hardness (Fig. 3b); no significant drug-related AEs |
| 2 | 34/F | Painful SCH since age 7; recurrence after prior treatment | Multiple excisions; sirolimus (temporary improvement, recurrence) | 500 mg daily; >12 months | Rapid improvement: ↓pain; ↑mobility (Fig. 3d); continued improvement | No persistent side effects requiring dose adjustment |
| 3 | Male; age NR | Congenital RLE lymphedema; angiosarcoma at 19 (biopsy R lower leg); later Dabska tumors; painful progressive osseous lesions (T1, T9, R 9th rib) | Below-knee amputation; adjuvant chemotherapy (doxorubicin, ifosfamide, paclitaxel); broad-spectrum tyrosine kinase inhibitor (no improvement) | 500 mg daily; >2 years | Pain resolution; increased sclerosis consistent with response (Fig. 3e); no progression reported | Osseous lesions biopsy: atypical vascular lesions without angiosarcoma; mild intermittent grade 1 diarrhea, no dose adjustment required |

Abbreviations: SCH, spindle cell hemangioma; RLE, right lower extremity; TKI, tyrosine kinase inhibitor; AE, adverse event; NR, not reported.

Extended Data Table 3. Primer and synthetic DNA sequences

IDH1_blockseq_f: GGCGTCAAATGTGCCACTATC

IDH1_block: CCATAAGCATGACGACCTATGATGATAGGTTaatt

IDH1_blockseq_r: CATGACTTACTTGATCCCCATAAGCAT

IDH2_blockseq_f: CTGGTTGAAAGATGGCGGCT

IDH2_block: GCCATGGGCGTGCCTGCCAAacca

IDH2_blockseq_r: TACCTGGTCGCCATGGGC

IDH1_WT_gblock: tgtcaaggtttattggatccaccatgtccaaaaaaatcagtggcggttctgtggtagagatgcaaggagatgaaatgacacgaatcatttgggaattgattaaagagaaactcatttttccctacgtggaattggatctacatagctatgatttaggcatagagaatcgtgatgccaccaacgaccaagtcaccaaggatgctgcagaagctataaagaagcataatgttggcgtcaaatgtgccactatcactcctgatgagaagagggttgaggagttcaagttgaaacaaatgtggaaatcaccaaatggcaccatacgaaatattctgggtggcacggtcttcagagaagccattatctgcaaaaatatcccccggcttgtgagtggatgggtaaaacctatcatcataggtcgtcatgcttatggggatcaatacagagcaactgattttgttgttcctgggcctggaaaagtagagataacctacacaccaagtgacggaacccaaaaggtgacatacctggtacataactttgaagaaggtggtggtgttgccatggggatgtataatcaagataagtcaattgaagattttgcacacagttccttccaaatggctctgtctaagggttggcctttgtatctgagcaccaaaaacactattctgaagaaatatgatgggcgttttaaagacatctttcaggagatatatgacaagcagtacaagtcccagtttgaagctcaaaagatctggtatgagcataggctcatcgacgacatggtggcccaagctatgaaatcagagggaggcttcatctgggcctgtaaaaactatgatggtgacgtgcagtcggactctgtggcccaagggtatggctctctcggcatgatgaccagcgtgctggtttgtccagatggcaagacagtagaagcagaggctgcccacgggactgtaacccgtcactaccgcatgtaccagaaaggacaggagacgtccaccaatcccattgcttccatttttgcctggaccagagggttagcccacagagcaaagcttgataacaataaagagcttgccttctttgcaaatgctttggaagaagtctctattgagacaattgaggctggcttcatgaccaaggacttggctgcttgcattaaaggtttacccaatgtgcaacgttctgactacttgaatacatttgagttcatggataaacttggagaaaacttgaagatcaaactagctcaggccaaactttaagttcataccggtgctaagaaggataattgtct

IDH1_R132C_gblock: tgtcaaggtttattggatccaccatgtccaaaaaaatcagtggcggttctgtggtagagatgcaaggagatgaaatgacacgaatcatttgggaattgattaaagagaaactcatttttccctacgtggaattggatctacatagctatgatttaggcatagagaatcgtgatgccaccaacgaccaagtcaccaaggatgctgcagaagctataaagaagcataatgttggcgtcaaatgtgccactatcactcctgatgagaagagggttgaggagttcaagttgaaacaaatgtggaaatcaccaaatggcaccatacgaaatattctgggtggcacggtcttcagagaagccattatctgcaaaaatatcccccggcttgtgagtggatgggtaaaacctatcatcataggtTgtcatgcttatggggatcaatacagagcaactgattttgttgttcctgggcctggaaaagtagagataacctacacaccaagtgacggaacccaaaaggtgacatacctggtacataactttgaagaaggtggtggtgttgccatggggatgtataatcaagataagtcaattgaagattttgcacacagttccttccaaatggctctgtctaagggttggcctttgtatctgagcaccaaaaacactattctgaagaaatatgatgggcgttttaaagacatctttcaggagatatatgacaagcagtacaagtcccagtttgaagctcaaaagatctggtatgagcataggctcatcgacgacatggtggcccaagctatgaaatcagagggaggcttcatctgggcctgtaaaaactatgatggtgacgtgcagtcggactctgtggcccaagggtatggctctctcggcatgatgaccagcgtgctggtttgtccagatggcaagacagtagaagcagaggctgcccacgggactgtaacccgtcactaccgcatgtaccagaaaggacaggagacgtccaccaatcccattgcttccatttttgcctggaccagagggttagcccacagagcaaagcttgataacaataaagagcttgccttctttgcaaatgctttggaagaagtctctattgagacaattgaggctggcttcatgaccaaggacttggctgcttgcattaaaggtttacccaatgtgcaacgttctgactacttgaatacatttgagttcatggataaacttggagaaaacttgaagatcaaactagctcaggccaaactttaagttcataccggtgctaagaaggataattgtct

IDH1_R132H_gblock: tgtcaaggtttattggatccaccatgtccaaaaaaatcagtggcggttctgtggtagagatgcaaggagatgaaatgacacgaatcatttgggaattgattaaagagaaactcatttttccctacgtggaattggatctacatagctatgatttaggcatagagaatcgtgatgccaccaacgaccaagtcaccaaggatgctgcagaagctataaagaagcataatgttggcgtcaaatgtgccactatcactcctgatgagaagagggttgaggagttcaagttgaaacaaatgtggaaatcaccaaatggcaccatacgaaatattctgggtggcacggtcttcagagaagccattatctgcaaaaatatcccccggcttgtgagtggatgggtaaaacctatcatcataggtcAtcatgcttatggggatcaatacagagcaactgattttgttgttcctgggcctggaaaagtagagataacctacacaccaagtgacggaacccaaaaggtgacatacctggtacataactttgaagaaggtggtggtgttgccatggggatgtataatcaagataagtcaattgaagattttgcacacagttccttccaaatggctctgtctaagggttggcctttgtatctgagcaccaaaaacactattctgaagaaatatgatgggcgttttaaagacatctttcaggagatatatgacaagcagtacaagtcccagtttgaagctcaaaagatctggtatgagcataggctcatcgacgacatggtggcccaagctatgaaatcagagggaggcttcatctgggcctgtaaaaactatgatggtgacgtgcagtcggactctgtggcccaagggtatggctctctcggcatgatgaccagcgtgctggtttgtccagatggcaagacagtagaagcagaggctgcccacgggactgtaacccgtcactaccgcatgtaccagaaaggacaggagacgtccaccaatcccattgcttccatttttgcctggaccagagggttagcccacagagcaaagcttgataacaataaagagcttgccttctttgcaaatgctttggaagaagtctctattgagacaattgaggctggcttcatgaccaaggacttggctgcttgcattaaaggtttacccaatgtgcaacgttctgactacttgaatacatttgagttcatggataaacttggagaaaacttgaagatcaaactagctcaggccaaactttaagttcataccggtgctaagaaggataattgtct

IDH1_Bam_Koz_f: tattgGaTCcaCCatgtccaaaaaaatcagtggcgg

IDH_Sal_r: tagcGTCGACatgaacttaaagtttggcctgagctagtttg

VEGFA_f: TCACCAAGGCCAGCACATAG

VEGFA_r: AGGCCCACAGGGATTTTCTT

GLUT1_f: TGGCATCAACGCTGTCTTCT

GLUT1_r: CTAGCGCGATGGTCATGAGT

**Extended Data Figures**

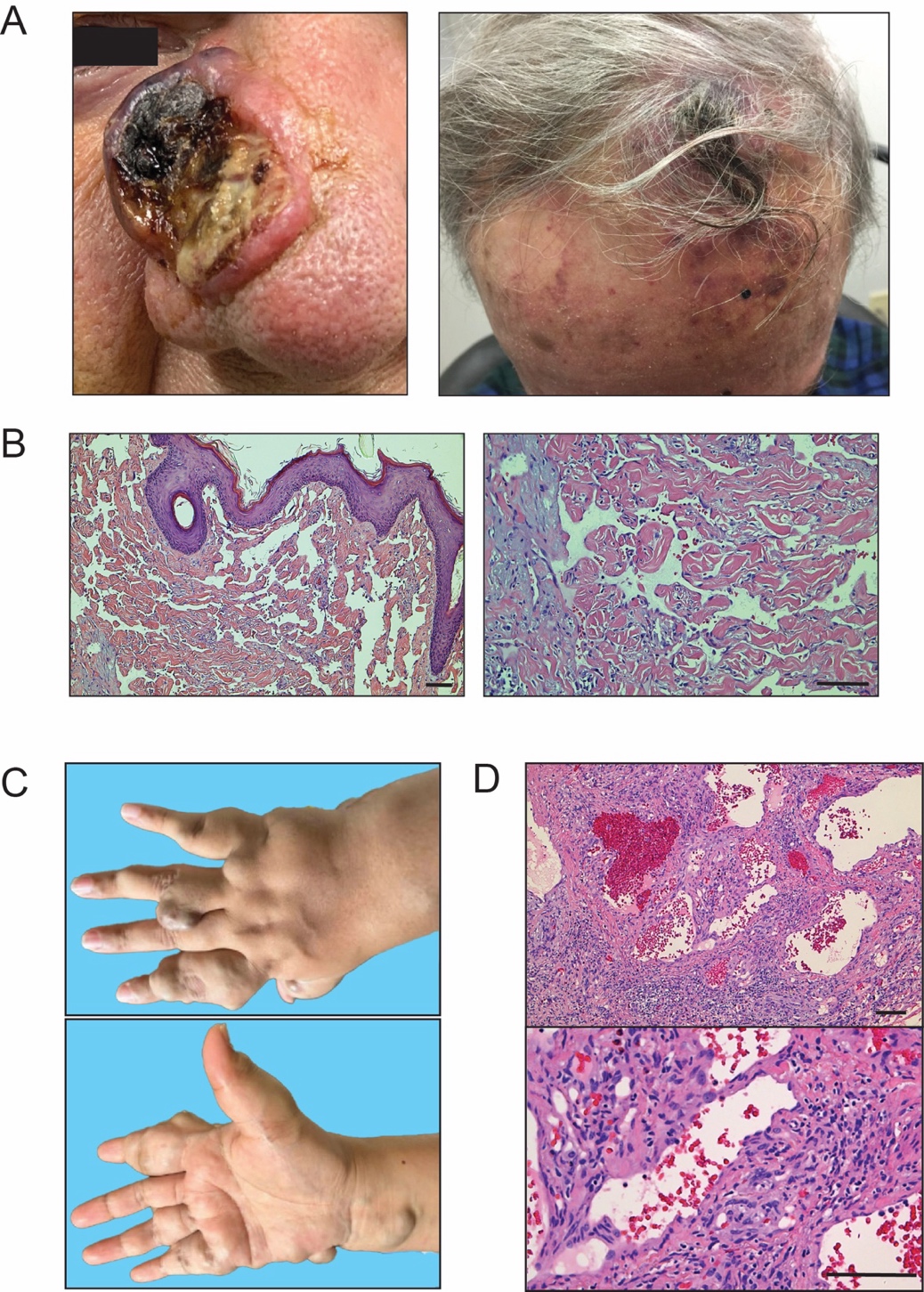

**Extended Data Figure 1.** **Clinical and histopathologic images of angiosarcoma and spindle cell hemangiomas (SCH).** (A) Representative clinical images of angiosarcoma, including an exophytic ulcerated nodule (left, CA2) and a violaceous, bruise-like patch (right, CA1). (B) Hematoxylin and eosin (H&E) staining of angiosarcoma (CA1) reveals irregular anastomosing vascular spaces lined by pleomorphic endothelial cells. (C) The hand of a Maffucci syndrome patient reveals the presence of multiple SCH lesions. (D) H&E staining of SCH from Maffucci patient reveals a spindle cell proliferation with associated large, dilated vascular spaces. Scale Bar = 100μm.

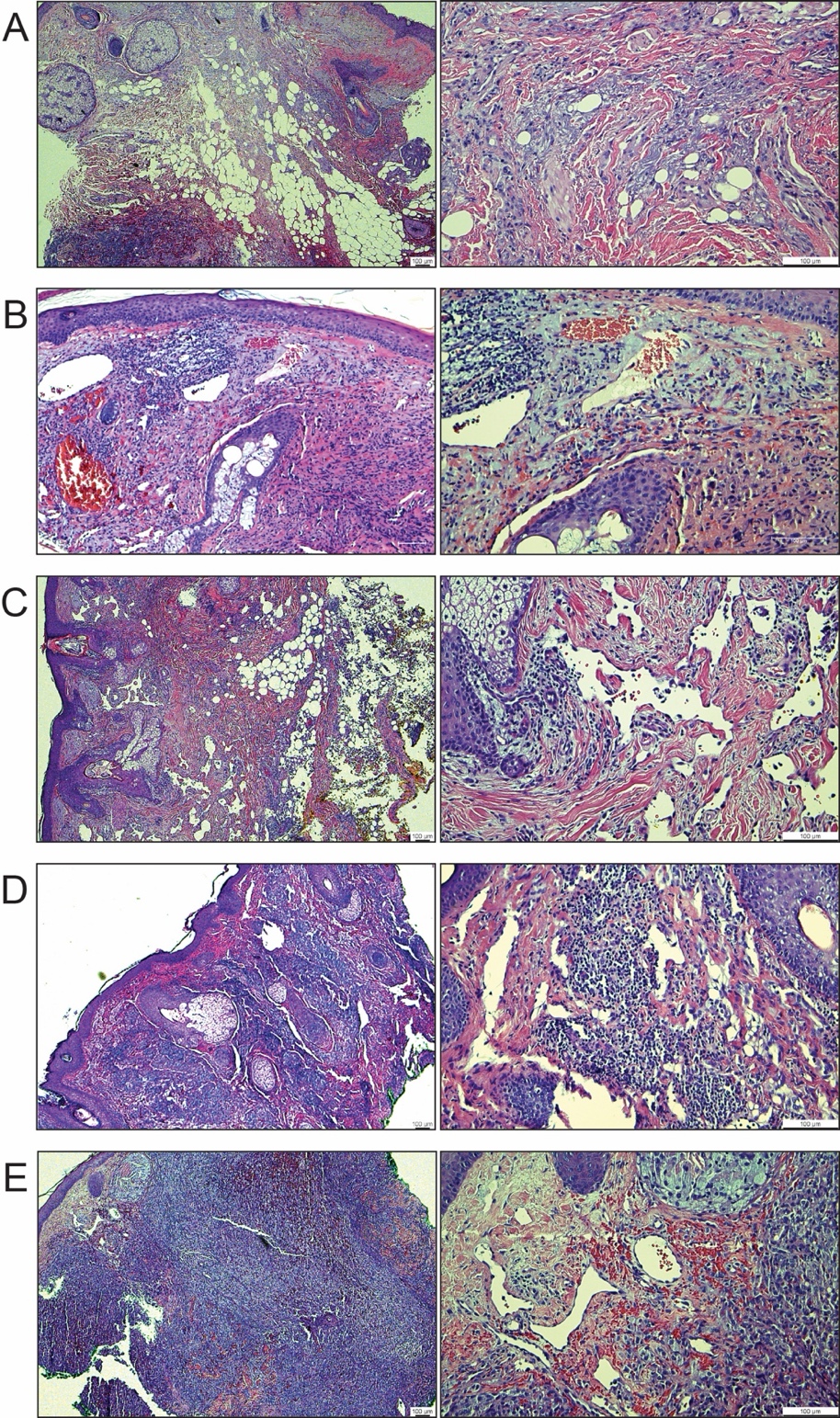

**Extended Data Figure 2. Additional H&E images of angiosarcoma.** Low power (5x) and high power (20x) images of CA2 (A), CA6 (B), CA17 (C), CA18 (D), and CA19 (E). Scale Bar = 100μm.

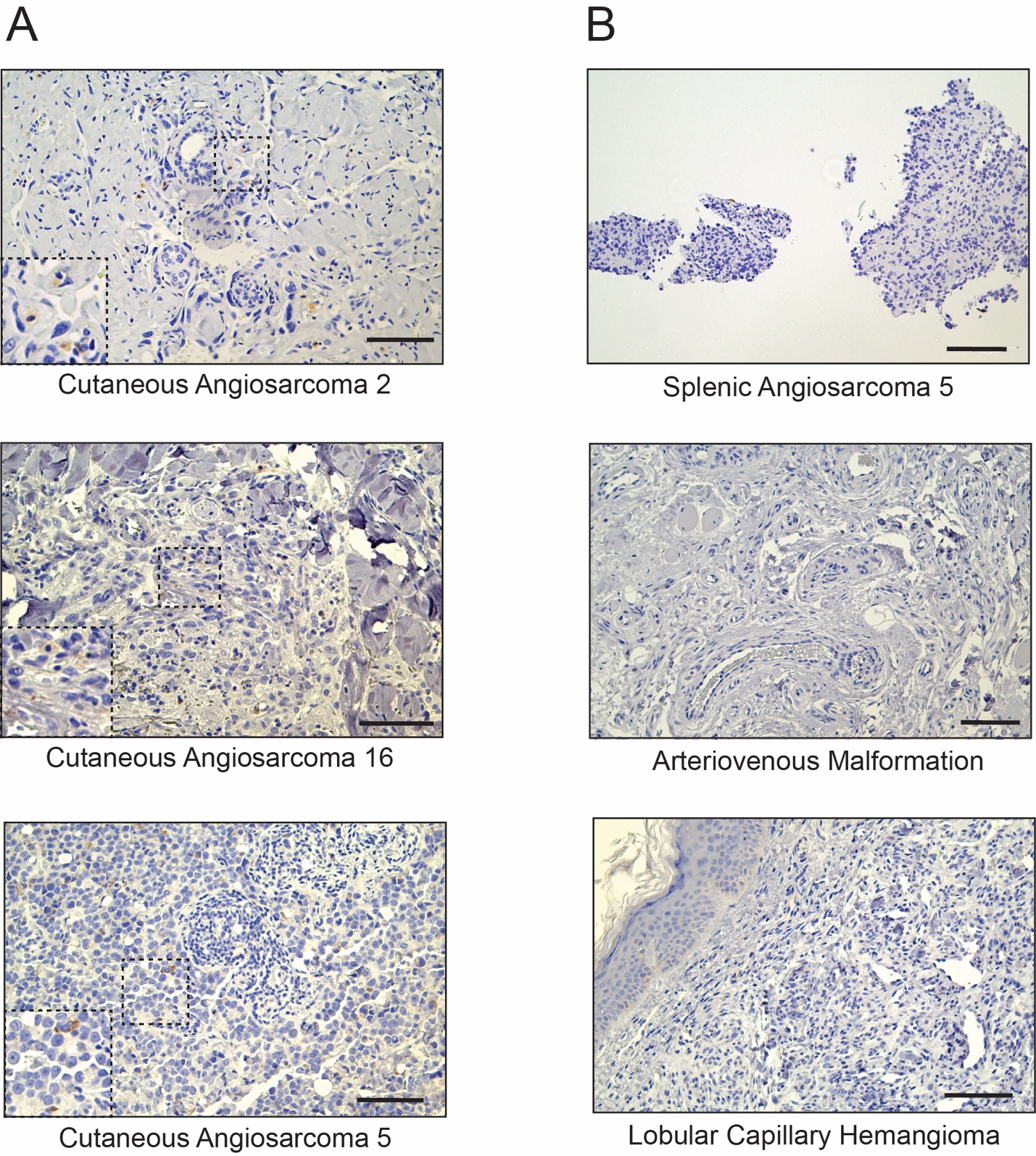

**Extended Data Figure 3. Additional IDH1 R132H IHC images.** (A) Immunohistochemistry (IHC) confirms cytoplasmic expression of IDH1 R132H in the indicated cutaneous angiosarcoma samples. (B) No expression of IDH1 R132H detected in the indicated splenic angiosarcoma and vascular controls.

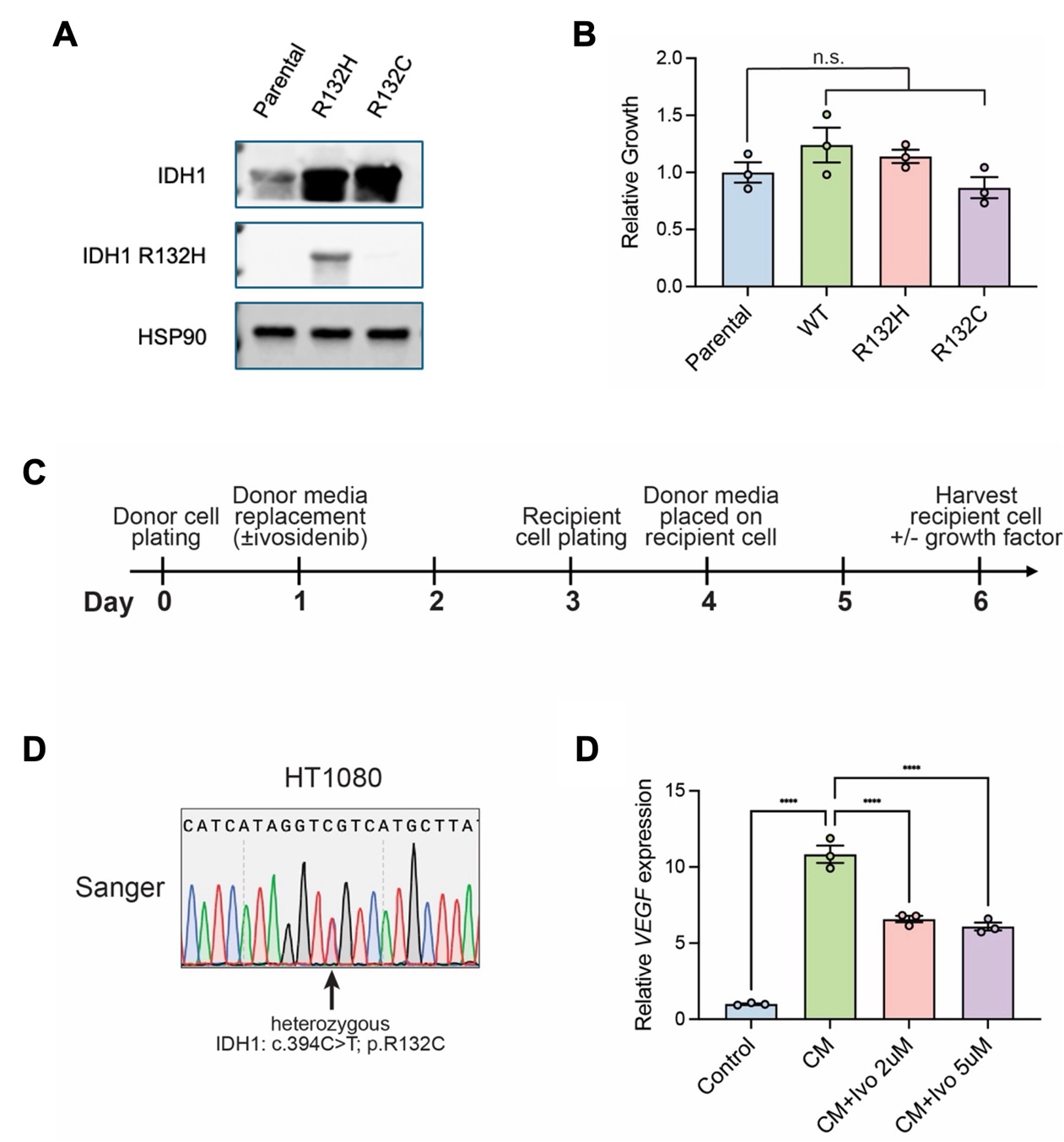

**Extended Data Figure 4. Effects of *IDH1* mutations on endothelial cells.** (A) Western blot confirms the expression of IDH1 R132C and R132H in immortalized human aortic endothelial cells (TeloHAECs). (B) Relative cell proliferation of wildtype (WT) or mutant IDH1 (R132H, R132C) in TeloHAECs as assessed by cell counts. (C) Schematic timeline of conditioned media (CM) experiments. (D) Sanger sequencing of *IDH1 c.394* in HT1080 fibrosarcoma cells demonstrating heterozygosity for IDH1 R132C. (E) qRT-PCR analysis of *VEGF* mRNA expression in TeloHAECs treated with CM derived from HT1080 cells for 48 h. CM induced VEGF expression, which was partially rescued by pre-treatment of HT1080 cells with ivosidenib (Ivo), 2 μM and 5 μM, as indicated. Bar graphs show mean ± s.e.m. from three independent biological replicates (n = 3). Statistical significance was determined using one-way ANOVA with Tukey’s multiple comparisons test. ****P < 0.0001.

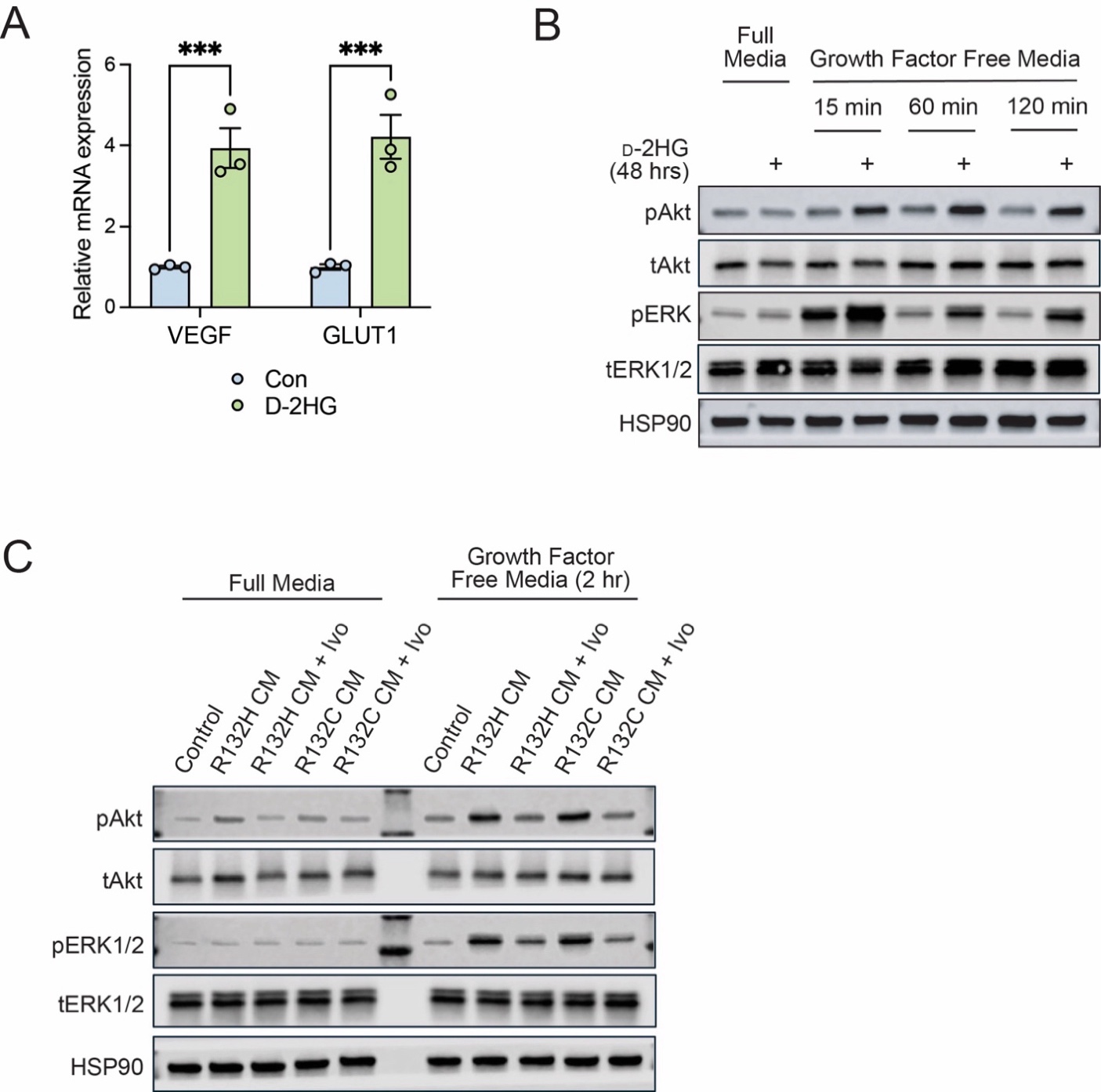

**Extended Data Figure 5. Conditioned media (CM) exerts non-cell autonomous effects on endothelial cells.** (A) Treatment of TeloHAEC cells with D-2-HG (800 μM) induced expression of *VEGF* and *GLUT1*. Data are presented as mean ± s.e.m. from independent biological replicates (n = 3 per group). Statistical significance was determined using unpaired two-tailed t-tests with correction for multiple comparisons. ***P < 0.001. (B) Western blot analysis showing time-dependent activation of pAkt and pERK1/2 signaling. TeloHAECs were incubated with D-2-HG (800 μM) for 48 h, followed by replacement with growth factor/serum-free media. Lysates were collected at 0, 15, 60, and 120 min after media replacement. (C) Western blot analysis of TeloHAECs incubated with CM from R132H or R132C mutant cells for 48 h, followed by media replacement with growth factor/serum-free media. 2 μM for 48 h. Indicated sample (+ Ivo) were treated with ivosidenib 2 μM throughout duration of treatment. Lysates were collected in regular endothelial cell media or 2 h after switch to growth factor-free media.

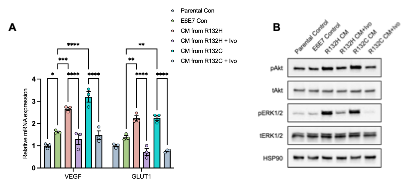

**Extended Data Figure 6.** IDH1 mutant alleles can exert non-cell autonomous effects on endothelial cells. (A) Relative mRNA expression of VEGF and SLC2A1 (GLUT1) in parental and E6E7-TeloHAECs following treatment with conditioned media (CM) from IDH1 R132H or R132C mutant cells, with or without pretreatment of donor cells with 2 μM ivosidenib (B) Immunoblot analysis of pAkt, total Akt, pERK1/2, total ERK1/2 and HSP90 in parental and E6E7 TeloHAECs treated with CM under the indicated conditions. Bar graphs show mean ± s.e.m. from three independent biological replicates (n = 3). Statistical analysis was performed using two-way ANOVA with Šídák’s multiple comparisons test. Exact P values are provided in the Source Data. *P < 0.05, **P < 0.01,***P < 0.001, ****P < 0.0001.

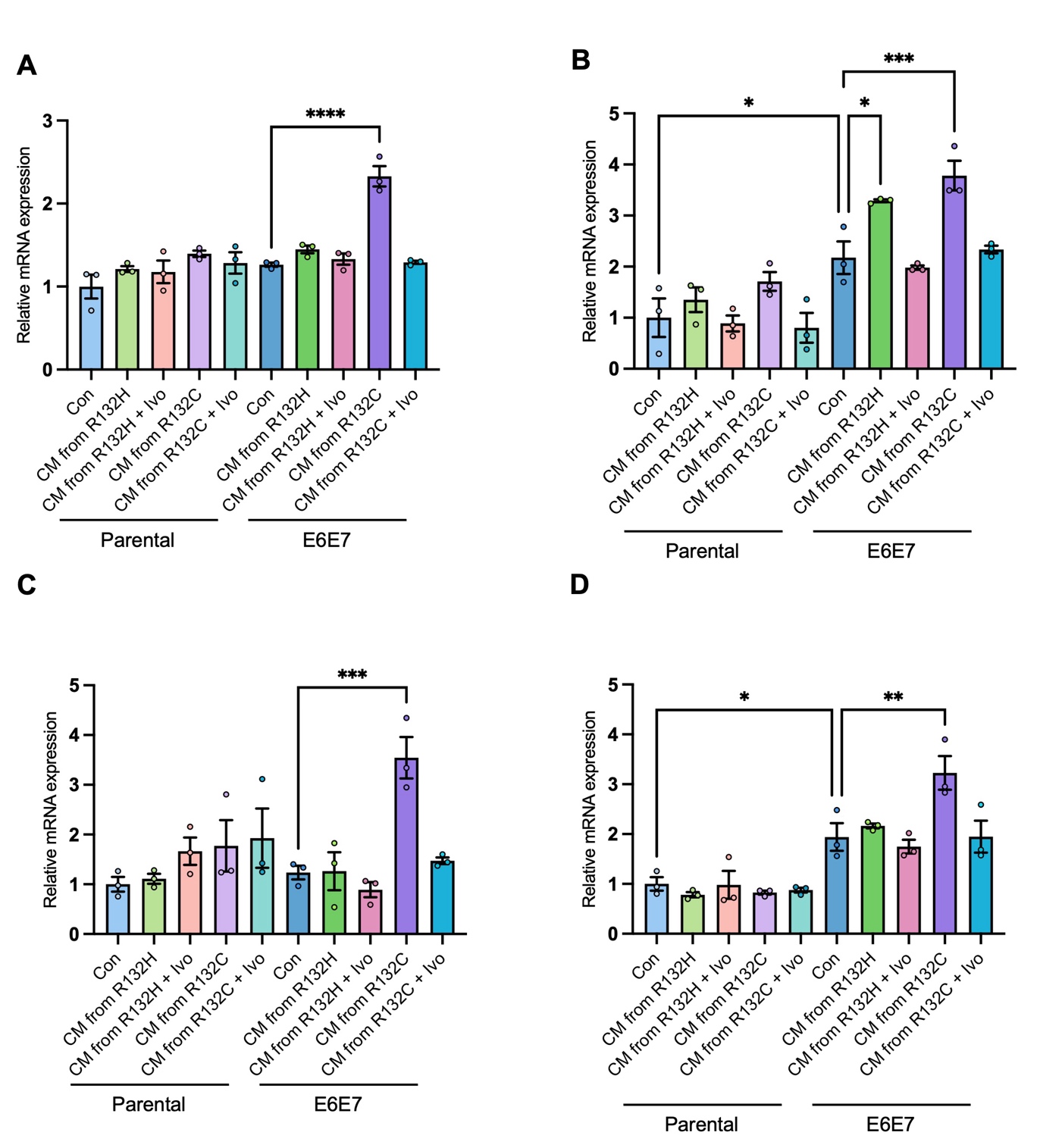

**Extended Data Figure 7.** IDH1 mutant conditioned media induces EMT-associated transcriptional changes in endothelial cells.(A–D) Relative mRNA expression of ZEB1 (A), SNAI1 (B), FSP1 (S100A4) (C), and VIM (D) in parental and E6E7 TeloHAECs following treatment with conditioned media (CM) from IDH1 R132H or R132C mutant cells, with or without pretreatment of donor cells with 2 μM ivosidenib.Bar graphs show mean ± s.e.m. from three independent biological replicates (n = 3). Statistical analysis was performed using two-way ANOVA with Šídák’s multiple comparisons test. Exact P values are provided in the Source Data. *P < 0.05.

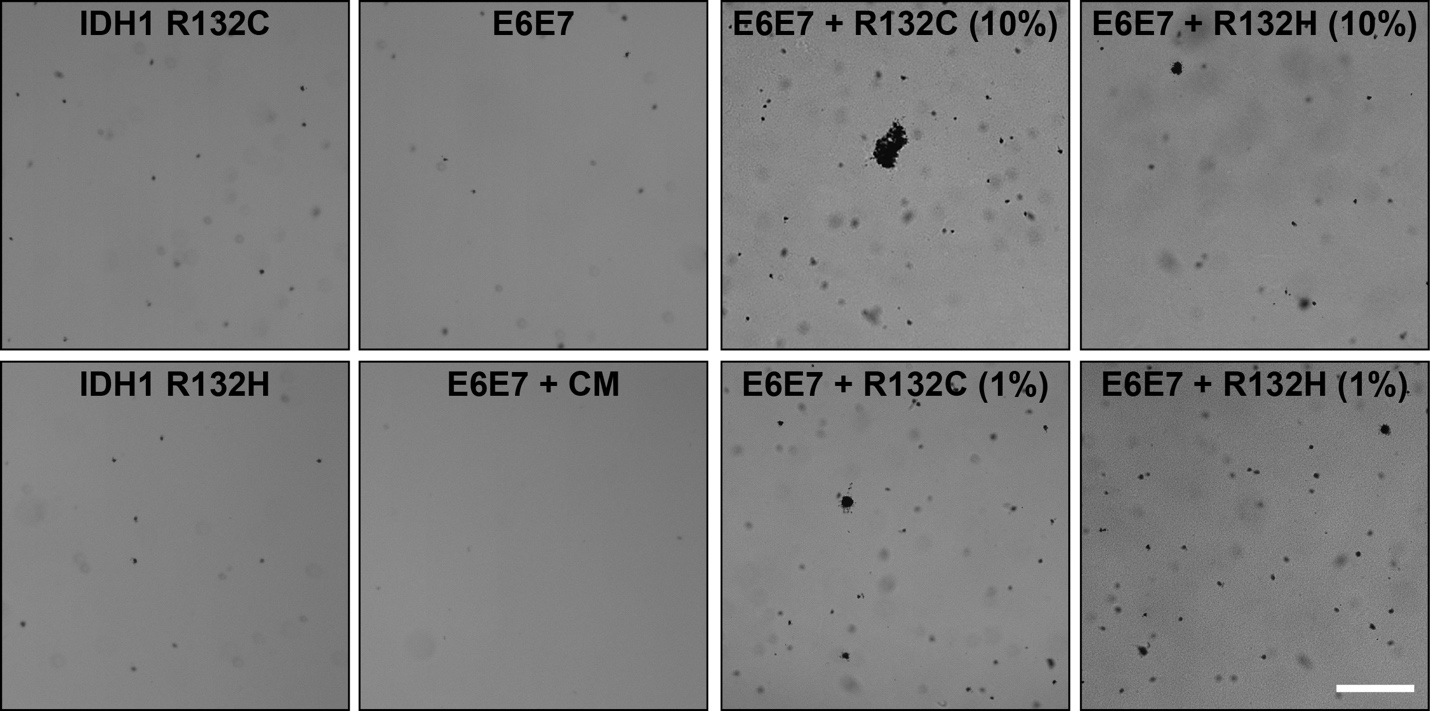

**Extended Data Figure 8.** Representative images of soft agar colony formation assays performed using TeloHAECs transduced with IDH1 mutant, E6E7, E6E7 treated with IDH1 R132C conditioned media (CM), or co-cultured with IDH1 mutant cells at the indicated ratios. Bar, 500 μm.
